# Variation in AMY2B Copy Number and Serum Amylase Activity in Wolves (*Canis lupus*), Brown Bears (*Ursus arctos*), and Red Foxes (*Vulpes vulpes*) from Bosnia and Herzegovina

**DOI:** 10.64898/2026.07.09.737415

**Authors:** Jasmin Katica, Ćazim Crnkić, Aida Kavazović, Dinaida Tahirović, Naris Pojskić, Vedad Škapur, Amira Koro-Spahić, Maja Varatanović, Teufik Goletić

## Abstract

The AMY2B gene encodes pancreatic amylase, a critical enzyme for starch digestion. While previous studies have examined AMY2B copy number variation (CNV) in domestic and some wild animals, less is known about wild carnivores inhabiting regions with limited anthropogenic starch exposure. We analyzed blood samples for serum amylase activity and copy number variation in AMY2B gene from 8 wolves (*Canis lupus*), 11 brown bears (*Ursus arctos*), and 3 red foxes (*Vulpes vulpes*) from Bosnia and Herzegovina. AMY2B gene copy number was assessed using droplet digital PCR (ddPCR), and serum amylase activity and glucose levels were quantified. Although the number of fox samples was limited, foxes and wolves consistently harbored two copies of AMY2B, while brown bears exhibited higher CNV (3.67–8.40, mean 5.88). Serum amylase activity was highest in foxes, moderate in wolves, and variable but lower in bears. Despite differences in AMY2B copy number and serum amylase activity, circulating glucose concentrations did not differ significantly among species. Our findings suggest that variation in AMY2B copy number among wild carnivores may be associated with species-specific evolutionary histories and dietary adaptations, providing insight into genomic mechanisms underlying carbohydrate utilization in natural populations.

## Introduction

The AMY2B gene encodes pancreatic amylase, a key enzyme involved in starch digestion, and variation in its copy number has been associated with dietary adaptation in several mammalian species. Despite increasing interest in AMY2B evolution, information on copy number variation in wild carnivore populations from the Dinaric–Balkan region remains limited. Gene copy number variation (CNV) itself is an important mechanism of evolutionary adaptation, enabling species to respond to environmental and dietary pressures. Increased AMY2B copy number in domestic dogs has been linked to adaptation to starch-rich diets accompanying domestication, whereas wild carnivores generally exhibit lower copy numbers consistent with their natural feeding ecology [1]. Grey wolves (*Canis lupus*), red foxes (*Vulpes vulpes*), and brown bears (*Ursus arctos*) differ substantially in their dietary habits. Wolves are predominantly carnivorous, while foxes and brown bears are opportunistic omnivores, consuming considerable amounts of plant-derived foods in addition to animal prey. These differences provide an opportunity to investigate whether variation in AMY2B copy number is associated with species-specific dietary ecology.

This study investigated AMY2B copy number variation in wolves, brown bears, and red foxes from Bosnia and Herzegovina using droplet digital PCR (ddPCR) and evaluated its association with serum amylase activity and glucose concentrations.

## Materials and methods

### Ethics statement

All procedures involving animals were conducted in accordance with national legislation and institutional guidelines governing animal welfare and wildlife research. The study protocol was approved by the Ethics Committee of the Veterinary Faculty, University of Sarajevo (Approval No. 07-03-461-2/26). Animal handling, chemical immobilization, blood collection, and recovery procedures were performed under the supervision of an experienced veterinarian, and all efforts were made to minimize stress, discomfort, and handling time.

### Sample collection

Blood samples were collected from 8 grey wolves (*Canis lupus*), 11 brown bears (*Ursus arctos*), and 3 red foxes (*Vulpes vulpes*) from Bosnia and Herzegovina. Of the wolves, two originated from free-ranging populations and six were maintained in captivity. Among the bears, five were free-ranging and six were captive. All foxes were captive individuals. Captive animals had originally been obtained from wild populations.

The wolf group consisted of three juveniles (2–3 months of age) and five adults, whereas the bear group included one juvenile (6 months of age) and ten adults. Blood samples were collected from the cephalic vein in wolves and foxes and from the femoral vein in bears. Wolves and bears were chemically immobilized under veterinary supervision using medetomidine–ketamine protocol and anesthesia was reversed with atipamezole (0.125 mg/kg in bears; 0.4 mg/kg in wolves). Foxes were manually restrained using humane capture devices and sampled without anesthesia.

### Anesthesia protocol

Chemical immobilization of wolves and bears was performed under the supervision of an experienced wildlife veterinarian. Following clinical assessment, animals were anesthetized using a medetomidine–ketamine combination administered remotely by dart gun. Bears received 0.025 mg/kg medetomidine and 5 mg/kg ketamine, whereas wolves received 0.08 mg/kg medetomidine and 4 mg/kg ketamine. Reversal of anesthesia was achieved using atipamezole at doses of 0.125 mg/kg and 0.4 mg/kg in bears and wolves, respectively. Animals remained under anesthesia only for the time required for blood collection and all individuals recovered without complications.

### Serum preparation and biochemical analyses

Blood samples were collected into EDTA tubes for DNA extraction and into serum separator tubes for biochemical analyses. Serum was separated by centrifugation and stored at −20°C until analysis. Serum amylase activity and glucose concentrations were determined using an IDEXX Catalyst One analyzer (IDEXX Laboratories, USA) employing dry-slide enzymatic methodology according to the manufacturer’s instructions. All blood samples were handled in accordance with standard biosafety procedures, including the use of personal protective equipment, appropriate sample labeling, and storage in leak-proof containers during transport and processing. Amylase activity was initially expressed as U/L and subsequently converted to µkat/L using the appropriate conversion factor.

### DNA extraction

Genomic DNA was extracted from EDTA-anticoagulated whole blood using the DNeasy Blood & Tissue Kit (Qiagen, Germany) according to the manufacturer’s instructions. DNA was eluted in 50 μL PCR-grade water. Initial DNA quantification was performed fluorometrically using the Qubit 1X dsDNA HS Assay Kit, yielding concentrations of 9–20 ng/μL. DNA concentration and purity were subsequently reassessed using a NanoDrop spectrophotometer (Thermo Fisher Scientific, USA) prior to ddPCR analysis. Samples with DNA concentrations between 5 and 40 ng/μL were considered suitable for downstream analyses and included in the study.

### Primer and probe design

Primer and probe sequences used for the AMY2B target in wolves and red foxes were adopted from a previously published ddPCR assay developed for domestic dogs [2]. Because this assay was originally validated in dogs, its application to other canid species should be interpreted in the context of the expected conservation of the target region among closely related taxa and previous evidence of AMY2B copy number variation across canids.

For brown bears, a separate ursid-specific assay was designed because a previously validated brown-bear-specific AMY2B ddPCR assay was not available. The canine AMY2B and C7orf28b-3 amplicon sequences were used as query sequences in BLAST searches against the available genomes of polar bear (Ursus maritimus) and American black bear (Ursus americanus). Homologous regions corresponding to a pancreatic amylase gene target (AMY/AMY2B-like) and to the putative single-copy reference gene CCZ1 (coiled-coil and C2 domain containing 1; formerly annotated as C7orf28b-3) were identified in both ursid genomes. CCZ1 was selected as the reference gene to maintain consistency with the previously validated canine ddPCR assay and because it represents a stable single-copy locus suitable for copy number normalization. Sequence comparison showed conservation of the selected target and reference regions between the queried ursid genomes. These conserved regions were subsequently used for primer and hydrolysis probe design using AlleleID software (Premier Biosoft, USA).

The resulting primer and probe sequences for the bear pancreatic amylase target were: AMY_Fw (5′-GACACTACCGAAATATCTGTTAATG-3′), AMY_Rv (5′- CGTGGAACGCACATAGTC-3′), and AMY_Pro (FAM- CGCAAGATCAAGAAGACCAACCAG-BHQ), generating a 172 bp amplicon. For the reference gene CCZ1, primers and probe were: CCZ1_Fw (5′-GGAGGTTGATGGTGCTAGG-3′), CCZ1_Rv (5′-GCCAACGCATCCCTTCTC-3′), and CCZ1_Pro (HEX-TCTCACTGTCACCAACCTGCCG-BHQ), generating a 165 bp amplicon. All oligonucleotides were synthesized by Generi Biotech (Czech Republic), with primers supplied desalted and probes HPLC-purified.

Assay optimization for the bear-specific ddPCR assay was performed using the QX200 Droplet Digital PCR platform (Bio-Rad, USA). Based on optimization experiments, primer and probe concentrations were adjusted to 900 nM and 500 nM, respectively, for both the amylase target and the CCZ1 reference assay. The optimized annealing/extension temperature for the bear-specific assay was 57°C, which provided improved droplet cluster separation and assay performance.

### Droplet digital PCR analysis

Copy number estimates for the amylase target were obtained using droplet digital PCR (ddPCR) on a QX200 Droplet Digital PCR System (Bio-Rad, USA), using a protocol adapted from Axelsson et al. [1]. To improve target accessibility and reduce DNA complexity before droplet generation, genomic DNA was digested with BamHI (New England Biolabs, USA). Copy number estimates obtained after restriction digestion were used as the final values reported in this study.

For restriction digestion, 5 µL of genomic DNA was incubated in a 10 µL reaction containing 1 U/µL BamHI enzyme and CutSmart buffer at 37°C for 60 min. Subsequently, 2 µL of digested DNA was used in each 20 µL ddPCR reaction.

Droplets were generated using a QX200 Droplet Generator according to the manufacturer’s instructions, yielding approximately 11,000 droplets per reaction (range: 9,000–15,000). Amplification was performed on a C1000 Touch thermal cycler (Bio-Rad) with an initial enzyme activation step at 95°C for 10 min, followed by 40 cycles of 95°C for 30 s and an annealing/extension step for 60 s, and a final droplet stabilisation step at 98°C for 10 min followed by cooling to 4°C. For wolves and red foxes, the annealing/extension temperature was 60°C and reactions contained 450 nM primers and 250 nM probes. For the bear-specific assay, the annealing/extension temperature was adjusted to 57°C based on assay optimisation, and reactions contained 900 nM primers and 500 nM probes to improve droplet cluster separation.

Following amplification, droplets were analysed using a QX200 Droplet Reader and QuantaSoft software version 1.6.6.0320 (Bio-Rad). Copy number estimates were calculated as the ratio between the concentration of the amylase target and the CCZ1 reference assay, multiplied by two to account for the diploid state of the reference locus. Because ddPCR provides quantitative estimates rather than integer counts, values are reported as copy number estimates.

### Statistical analysis

One-way ANOVA was used to assess differences in the studied parameters among species, followed by Tukey’s post hoc test when a significant species effect was detected. Regression analyses and Pearson’s correlation coefficients were applied to evaluate associations between CNV and enzyme activity. Differences between two groups (wilderness vs. captivity; young vs. adult animals) with small sample sizes (n < 5) were analyzed using the nonparametric Mann–Whitney test. A P value < 0.05 was considered statistically significant. Results are presented graphically or in tables as means (X̄) with appropriate measures of variability, or as medians (X̃) when nonparametric tests were applied. All analyses were conducted using Minitab 17.

## Results

Copy number estimates for the AMY2B target in canids were highly consistent, averaging 2.0 ± 0.05 in wolves and 1.9 ± 0.08 in red foxes (Fig. 1). In bears, copy number estimates for the pancreatic amylase (AMY/AMY2B-like) target were higher, averaging 5.9 ± 1.4 and ranging from 3.7 to 8.4. All examined bears except one had copy number estimates above four copies.

**Figure 1.**
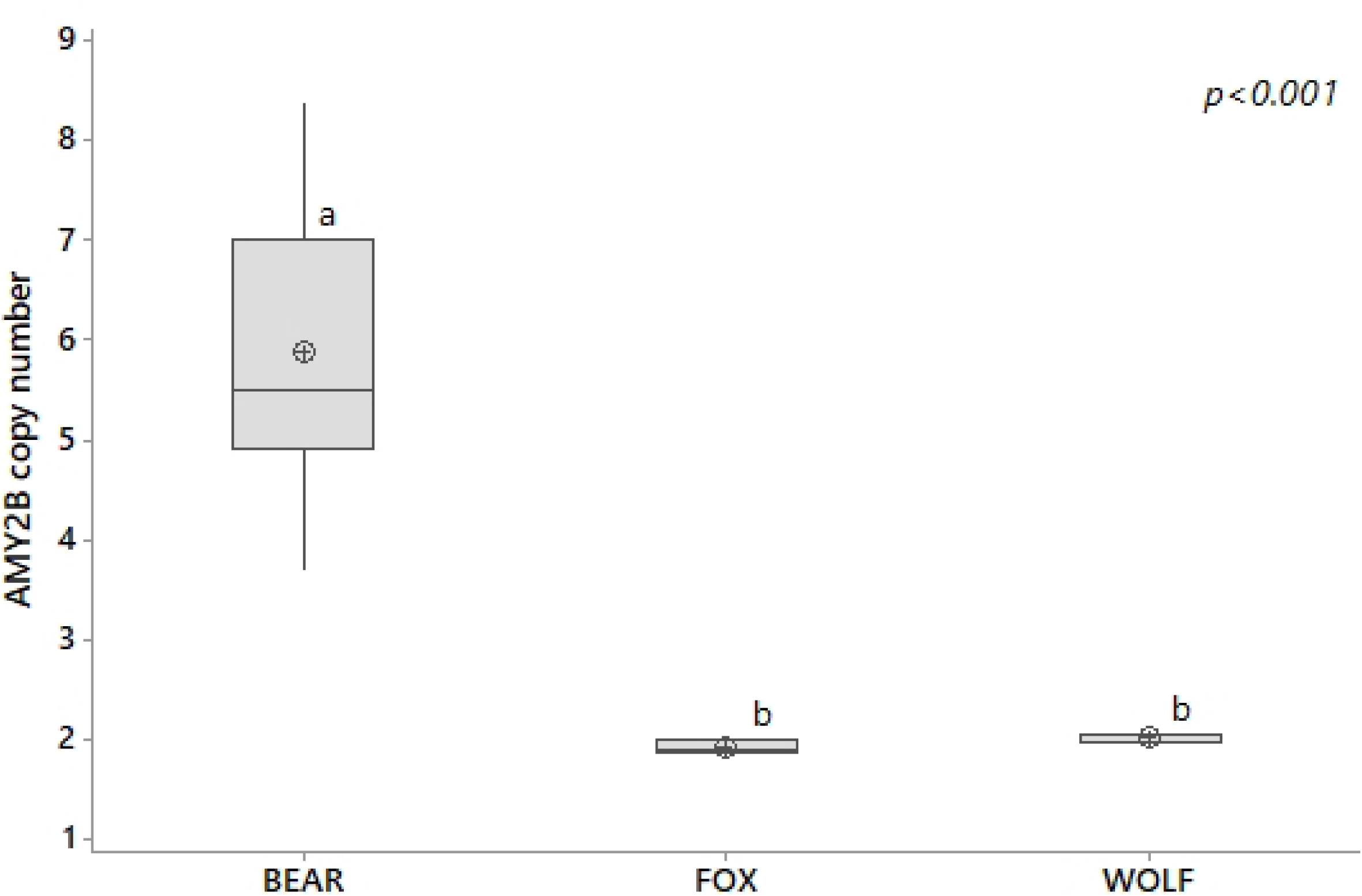
Copy number of *AMY2B* gene in wild carnivores. ^ab^ - Different superscript letters indicate statistical significance (p<0.05) of differences between means.

Serum amylase activity showed an inverse interspecific pattern relative to amylase gene copy number estimates (Fig. 2 compared with Fig. 1). The lowest activity was observed in bears (1.2 ± 0.7), followed by wolves (3.6 ± 1.5) and red foxes (6.2 ± 0.9), with a statistically significant interspecific difference (p < 0.001).

**Figure 2.**
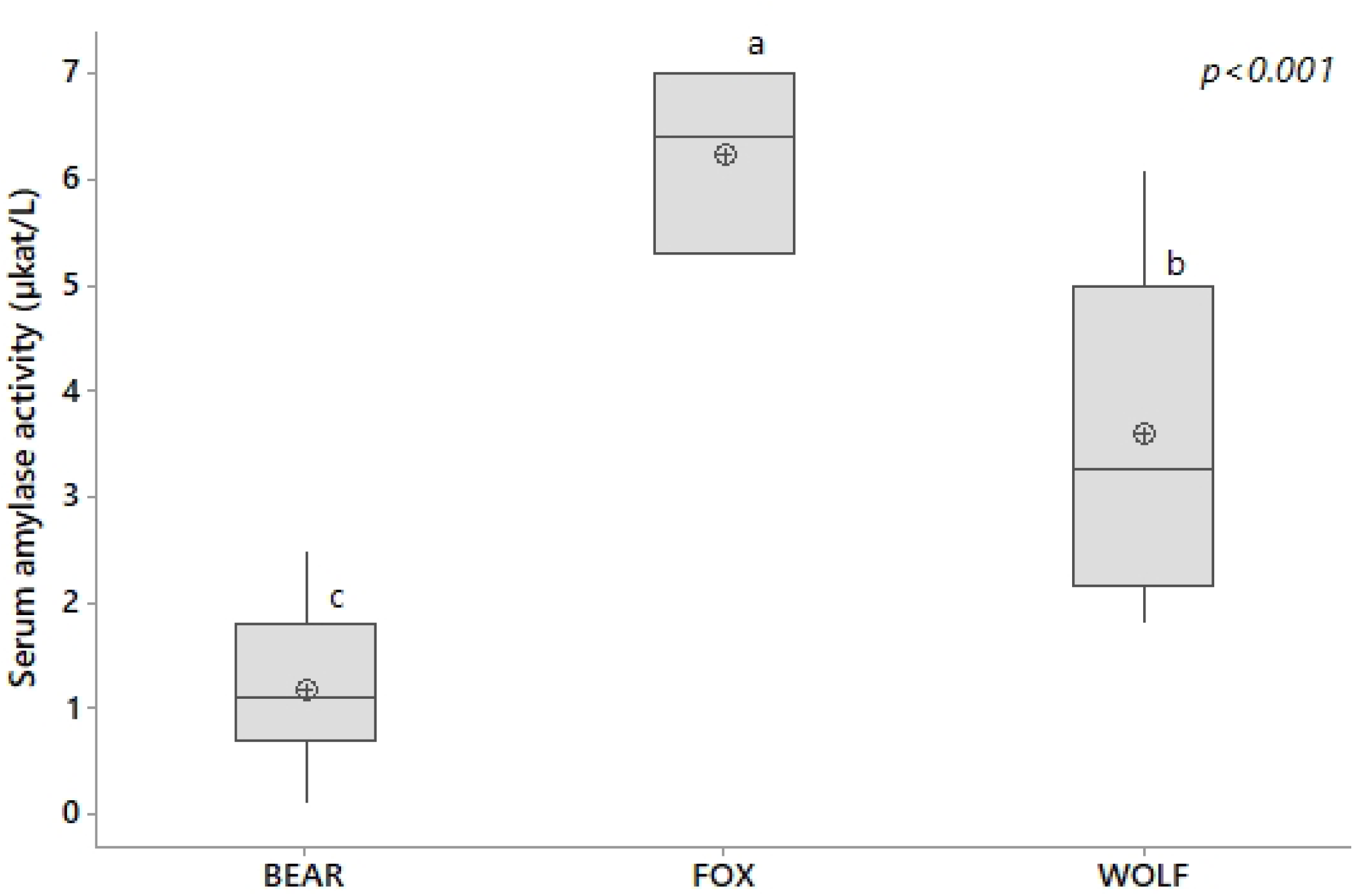
Serum amylase activity in wild carnivores. ^abc^ - Different superscript letters indicate statistical significance (p<0.05) of differences between means.

Serum glucose concentrations varied widely and showed numerical differences among species without reaching statistical significance (Fig. 3). Red foxes exhibited the highest mean glucose value (7.9 ± 2.2 mmol/L). In wolves and bears, serum glucose concentrations were 6.0 ± 1.5 and 6.1 ± 3.1 mmol/L, respectively. The widest range of serum glucose variation was observed in bears.

**Figure 3.**
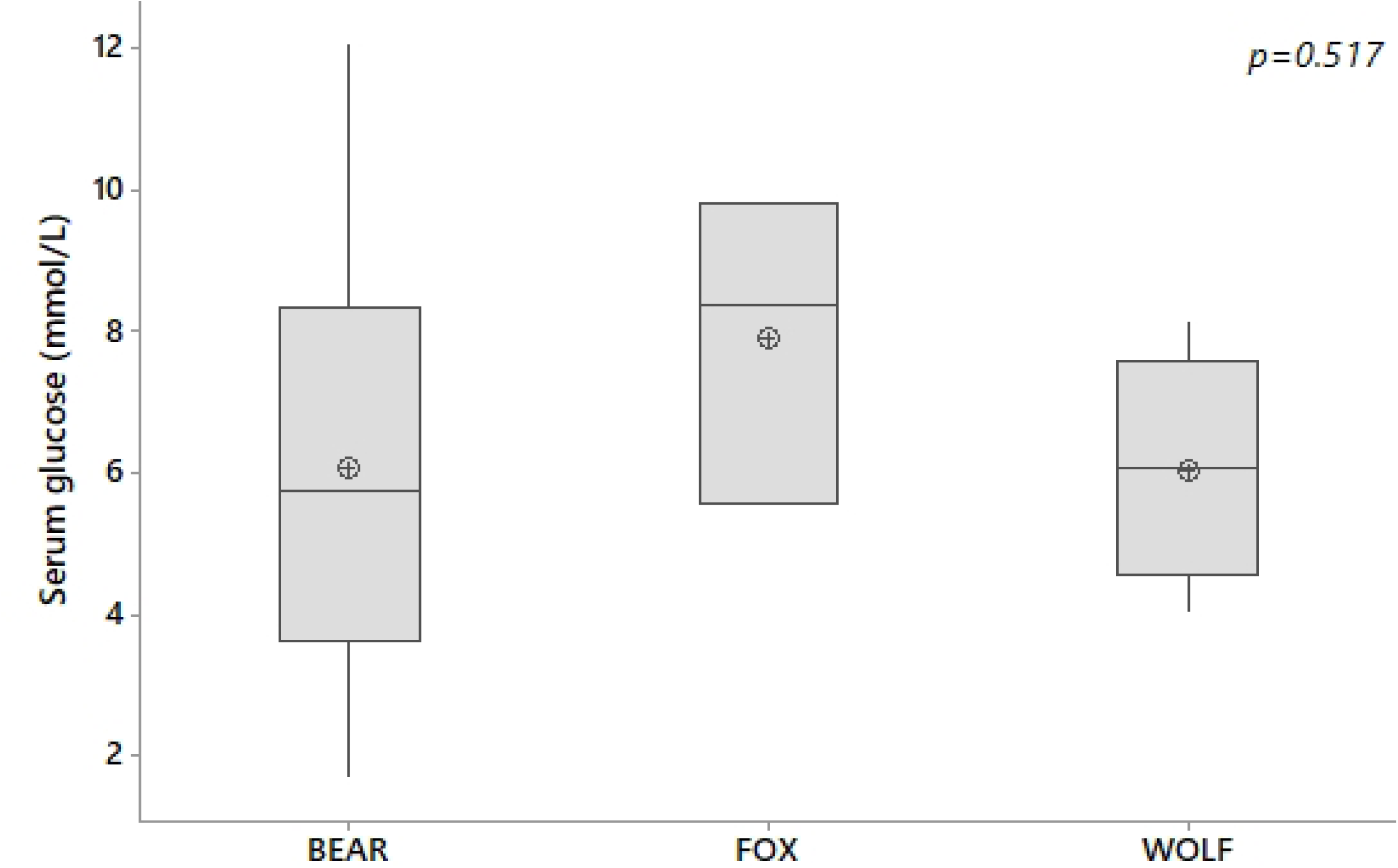
Serum glucose content in wild carnivores.

No statistically significant association was detected between amylase target copy number estimates and serum amylase activity, either in bears (r = 0.352; p = 0.289) or in canids (r = −0.534; p = 0.090). Therefore, the present data (Fig 4; Fig 5) do not support a direct relationship between copy number estimates and circulating amylase activity in the analyzed animals.

**Figure 4.**
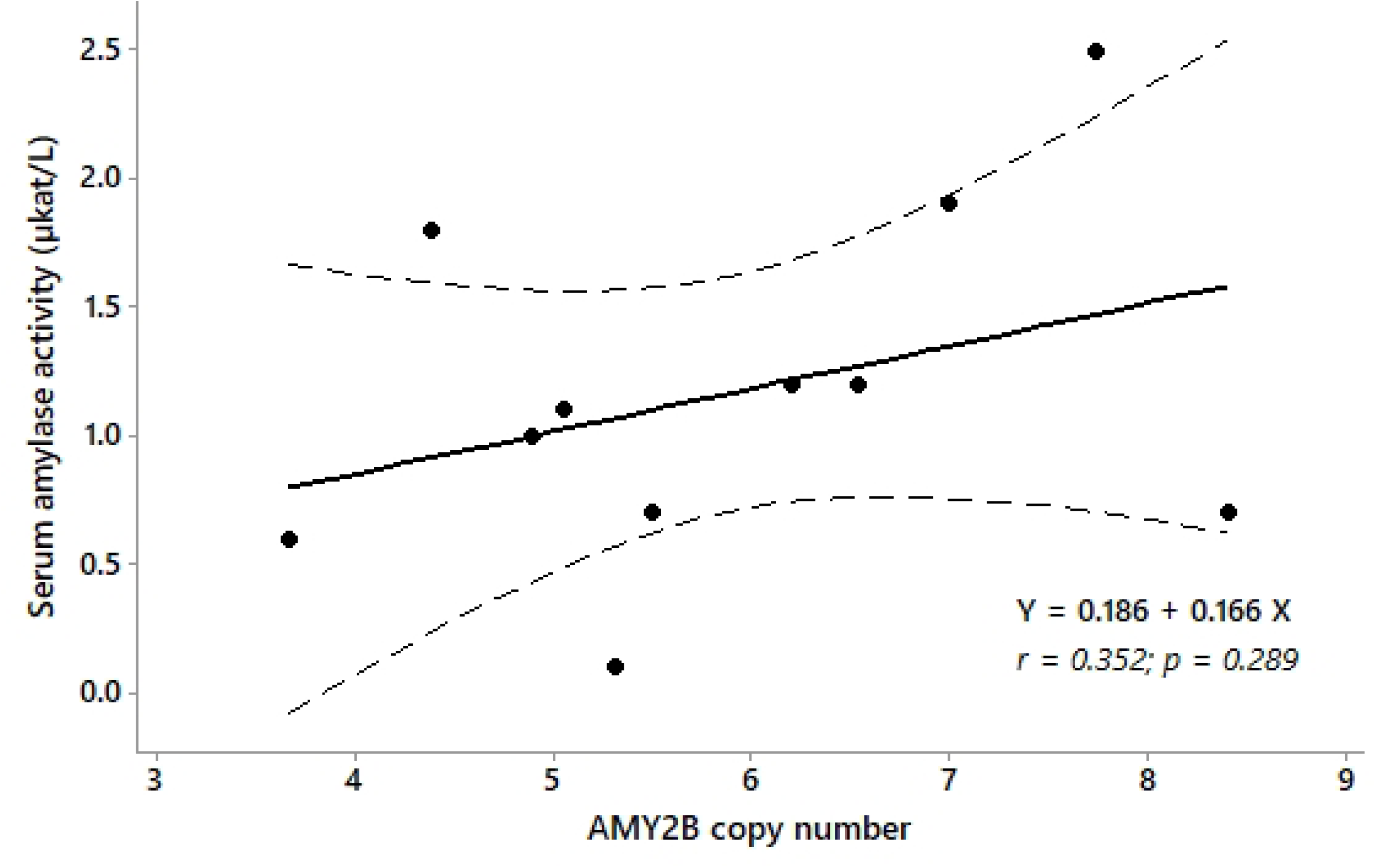
The relationship between serum amylase activity and AMY2B gene copy number in bears.

**Figure 5.**
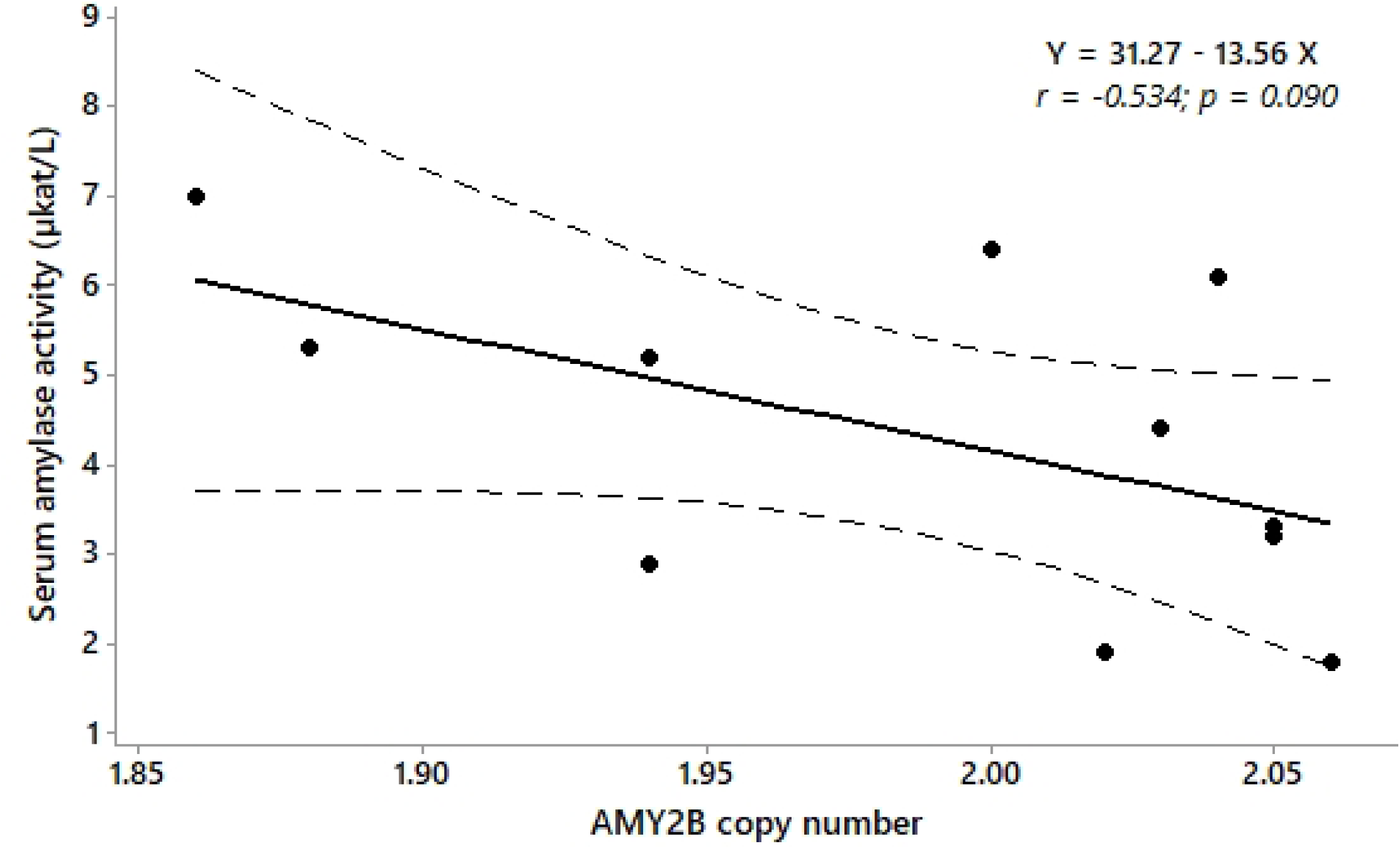
The relationship between serum amylase activity and AMY2B gene copy number in wild canids (wolves and foxes)

Two wolves and six bears originated from the wild, whereas the remaining wolves and bears were from captivity. Due to the relatively small group sizes, differences between these groups were assessed using the nonparametric Mann–Whitney test (Table 1). None of the investigated parameters differed significantly between wild and captive animals. All foxes included in the study originated from captivity.

**Table 1.** Median values (X̃) of AMY2B gene copy number, amylase activity, and serum glucose concentration in wild and captive bears and wolves.

|  | Species | Wild animals | Captive animals | p = |
| --- | --- | --- | --- | --- |
| AMY2B copy number | bear | 5.50 | 5.63 | 0.927 |
|  | wolf | 2.04 | 2.04 | 0.613 |
| Amylase ($\mu\text{kat/L}$) | bear | 0.70 | 1.15 | 0.714 |
|  | wolf | 1.85 | 3.85 | 0.067 |
| Glucose (mmol/L) | bear | 6.82 | 5.46 | 0.648 |
|  | wolf | 5.66 | 6.28 | 0.868 |

Of the eight wolves included in the study, three were pups aged 2-3 months and five were adults. The number of AMY2B gene copies, serum amylase activity, and serum glucose concentrations did not differ significantly between these age groups (Table 2). In the remaining two species, this comparison was not performed due to the insufficient number of individuals (bears) or the absence of young animals available for study (foxes).

**Table 2.** Median values (X̃) of AMY2B gene copy number, amylase activity, and serum glucose concentration in adult wolves and wolf pups.

|  | adult wolves | wolf pups | p = |
| --- | --- | --- | --- |
| AMY2B copy number | 2.04 | 2.03 | 0.763 |
| Amylase ($\mu\text{kat/L}$) | 3.30 | 3.20 | 1.000 |
| Glucose ($\text{mmol/L}$) | 5.70 | 6.86 | 0.766 |

## Discussion

### AMY2B copy number variation among wild carnivores

Copy number variation of amylase genes represents one of the most extensively studied examples of dietary adaptation in mammals. Increased copy numbers of pancreatic amylase genes have been associated with enhanced starch digestion capacity in several species and are often interpreted as genomic signatures of adaptation to distinct nutritional environments [1,3]. In domestic dogs and ancient European dog populations, expansion of the AMY2B gene has been linked to increased access to starch-rich food resources associated with domestication and close association with humans [1,4].

Dietary ecology of wild carnivores in Bosnia and Herzegovina may differ from that reported in more remote or less anthropogenically influenced populations. Bears and wolves increasingly utilise human-modified landscapes, particularly in areas where natural habitats overlap with agricultural land, livestock production, orchards, crop fields, waste disposal sites, and peri-urban settlements. Such environments may provide access to carbohydrate-rich food resources that are uncommon in strictly natural ecosystems. Consequently, dietary exposure to starch-rich foods may vary considerably among individuals and populations.

In contrast, captive animals are typically maintained on controlled diets with predictable nutritional composition and regular food availability. Depending on management practices, captive diets may contain either lower or higher proportions of carbohydrates than those consumed by free-ranging individuals. Therefore, comparisons between captive and wild populations should be interpreted cautiously, as differences in feeding ecology, seasonal food availability, and human-associated food resources may influence metabolic traits and potentially affect selective pressures acting on genes involved in carbohydrate digestion.

The present study revealed marked interspecific differences in amylase gene copy number among carnivores from Bosnia and Herzegovina. Brown bears exhibited substantially higher copy number estimates for the AMY/AMY2B-like target than wolves or red foxes, whereas both canid species carried approximately two copies of AMY2B. These findings are consistent with previously reported patterns in carnivorous and omnivorous mammals and support the hypothesis that long-term dietary ecology may have contributed to the evolutionary diversification of starch-digestion genes [3].

The elevated copy number estimates observed in brown bears are noteworthy in the context of their omnivorous diet. Unlike wolves, which are predominantly hypercarnivorous, brown bears regularly consume substantial quantities of fruits, berries, roots, and other plant-derived foods throughout much of the year. Such dietary flexibility has been considered a major ecological adaptation of bears and may have favored the maintenance or expansion of amylase gene copy number during their evolutionary history [3].

In contrast, the consistently low AMY2B copy number observed in wolves and red foxes suggests a more conserved genomic architecture in these canids. The low AMY2B copy number in red foxes is in agreement with previous ddPCR analyses demonstrating that most individuals possess two copies of the AMY2B gene [5].

Comparison with previously published data showed no statistically significant differences between wolves from Bosnia and Herzegovina and North American wolf populations reported by Pajic et al. [3]. This apparent conservation of AMY2B copy number across geographically distant wolf populations may reflect similar evolutionary constraints associated with a predominantly carnivorous lifestyle. Nevertheless, the slightly lower values observed in our study indicate that population-level variation cannot be excluded and should be explored using larger sample sizes and broader geographic sampling.

### Relationship between AMY2B copy number and serum amylase activity

Although brown bears exhibited the highest amylase target copy number estimates, they did not demonstrate the highest serum amylase activity. In contrast, red foxes displayed the highest amylase activity despite possessing only two copies of the AMY2B gene. Furthermore, no statistically significant association was detected between copy number estimates and circulating amylase activity.

These findings suggest that amylase gene copy number estimates alone are insufficient to explain variation in serum amylase activity and emphasize the complexity of genotype–phenotype relationships in nutritional adaptation. While gene duplication may increase the potential for enhanced enzyme production, actual enzyme activity is influenced by multiple regulatory mechanisms, including transcriptional regulation, tissue-specific gene expression, physiological status, environmental influences, and post-transcriptional processes. Similar observations have been reported in domestic dogs, where variation in AMY2B copy number was not consistently associated with serum amylase activity, indicating that additional regulatory mechanisms contribute to the regulation of carbohydrate metabolism [2].

The absence of a clear relationship between copy number estimates and enzyme activity is particularly important because it highlights the need to distinguish between genomic potential and realized physiological function. Consequently, caution should be exercised when interpreting amylase gene copy number as a direct proxy for digestive enzyme activity in wild populations.

### Comparison with previous biochemical studies

Biochemical parameters obtained for brown bears were compared with values previously reported for the Dinaric bear population by Huber et al. [6]. Glucose concentrations observed in the present study were generally consistent with previously published data, indicating similar patterns of glucose regulation across populations.

In contrast, mean serum amylase activity was higher than values reported previously. Several factors may contribute to this discrepancy, including differences in analytical methodologies, physiological status of sampled animals, environmental conditions, seasonal variation, and the inclusion of both captive and free-ranging individuals. Because detailed dietary information was unavailable for individual animals, the contribution of nutritional factors cannot be assessed directly and remains an important area for future investigation.

### Glucose homeostasis and metabolic adaptation

Despite substantial differences in amylase gene copy number estimates and serum amylase activity, glucose concentrations were remarkably similar among the studied species. This observation suggests that distinct evolutionary and physiological strategies may ultimately achieve comparable metabolic outcomes.

Glucose homeostasis in mammals is maintained through complex regulatory networks that extend far beyond starch digestion. Brown bears exhibit profound seasonal metabolic adaptations associated with hibernation, including reversible insulin resistance and altered carbohydrate metabolism [7]. In contrast, wolves and foxes rely predominantly on gluconeogenesis from proteins and lipids to maintain circulating glucose concentrations, consistent with the metabolic physiology of carnivorous mammals [8,9,10]. The convergence of glucose values observed in the present study likely reflects the effectiveness of these species-specific metabolic strategies rather than direct effects of amylase gene copy number variation.

Interestingly, these findings indicate that genomic differences affecting carbohydrate digestion do not necessarily translate into measurable differences in circulating glucose concentrations under physiological conditions. This observation further emphasizes the importance of integrating genomic and physiological data when investigating nutritional adaptation.

### Study limitations

Several limitations should be considered when interpreting the present findings. First, the number of analyzed animals was relatively small; particular caution is warranted when interpreting results for red foxes due to the very small sample size (n = 3). Second, both captive and free-ranging individuals were included in the study, potentially introducing environmental and nutritional variability. Third, detailed dietary histories were not available for individual animals, preventing direct assessment of relationships between food composition and AMY2B copy number variation. Fourth, for brown bears, the ddPCR assay was designed against conserved ursid pancreatic amylase regions rather than a previously published brown-bear-specific AMY2B assay; therefore, the corresponding values should be interpreted conservatively as copy number estimates for a pancreatic amylase target. Finally, serum amylase activity may not accurately reflect pancreatic AMY2B expression, and future studies incorporating transcriptomic or proteomic approaches would provide valuable insight into the functional consequences of gene copy number variation.

Future studies incorporating larger sample sizes, dietary characterization, gene expression analyses, and additional wild populations will help clarify the evolutionary and functional significance of AMY2B variation in carnivorous and omnivorous mammals.

## Conclusion

This study provides novel data on amylase gene copy number variation in wild carnivores from Bosnia and Herzegovina and contributes to a broader understanding of the genetic basis of dietary adaptation in mammals. Clear interspecific differences were observed, with wolves and red foxes consistently exhibiting approximately two AMY2B copies, whereas brown bears displayed substantially higher copy number estimates for the pancreatic amylase target.

These findings are consistent with previously reported associations between amylase gene expansion and broader dietary niches, and are compatible with the hypothesis that long-term feeding ecology has influenced amylase gene copy number variation. However, despite marked differences in gene copy number among species, no significant association was detected between copy number estimates and serum amylase activity. This indicates that gene dosage alone does not fully explain variation in amylase activity and suggests the involvement of additional regulatory and physiological mechanisms. Taken together, our results highlight the complexity of nutritional adaptation and emphasize that genomic variation and metabolic phenotype are not necessarily directly coupled. The observed pattern of elevated amylase gene copy number estimates in brown bears, compared with the conserved AMY2B copy number found in wolves and red foxes, provides further evidence that amylase gene evolution may reflect long-term ecological and dietary specialization in wild mammals.

Given the limited sample size and the inclusion of both captive and free-ranging individuals, the present findings should be considered exploratory. Future studies incorporating larger populations, detailed dietary data, and gene expression analyses will be necessary to better understand the functional significance of AMY2B copy number variation and its role in mammalian nutritional adaptation.

